# Designing proteins with reduced T-cell epitopes through policy optimization

**DOI:** 10.1101/2025.09.27.678937

**Authors:** Manvitha Ponnapati, Sapna Sinha, Brian Lynch, Edward S. Boyden, Joseph Jacobson

## Abstract

Deep generative models for protein structure and sequence are increasingly used to design proteins with therapeutic and industrial applications, but the clinical success of designed therapeutics ultimately depends on their compatibility with the human immune system. Immune response is triggered by a cascade of molecular events, including proteasomal cleavage, peptide elution, and binding to the highly polymorphic Major Histocompatibility Complex (MHC) Class I molecules, yet prior approaches have generally modeled these processes in isolation or restricted attention to a limited set of alleles. In this work, we develop predictors for cleavage, elution, and binding affinity in the MHC Class I pathway, incorporating evidential deep learning to provide unified uncertainty estimates. We first perform supervised fine-tuning of a protein language model on human proteins, and then align the model through group relative policy optimization (GRPO) to reduce MHC Class I epitopes under a curriculum learning framework, in which the curriculum progressively increases the number of masked predicted epitopes and the number of alleles considered during training. This strategy enables the generation of protein candidates that are optimized for immune compatibility across diverse MHC Class I alleles while accounting for predictive uncertainty in modeling the immune response.

## 1 Introduction

Design of proteins with tailored function is fundamental to modern biotechnology, with applications in therapeutics, synthetic biology, and bio-manufacturing (Koh et al., 2025). Modern machine learning models for protein sequence and structure, such as AlphaFold (Jumper et al., 2021), ESM (Hayes et al., 2025; Lin et al., 2023), and Boltz (Passaro et al., 2025), have enabled computational pipelines that explore vast regions of sequence space and accelerate the discovery and design of novel therapeutics and industrial proteins. These advances have allowed protein designers to create proteins with unprecedented diversity beyond naturally occurring sequences (Ruffolo et al., 2024; Hayes et al., 2024), demonstrating that machine learning models can effectively represent the complex sequence–structure–function relationships of proteins. While they have enabled the discovery of sequences for novel folds, backbone designs, and binders (Koh et al., 2025), downstream optimization for properties such as immunogenicity (Jawa et al., 2020) and aggregation remains challenging due to both limited data and the inherent complexity of biological systems.

A key representation of this complexity is the immunogenicity of novel proteins. Immune response can be elicited by various pathways, including MHC Class I, MHC Class II, or alternative antigen-processing mechanisms, depending on the context and type of protein delivery (Arneth, 2025). In this work, we focus on the MHC Class I pathway, though the approach can readily be extended to other pathways. The MHC Class I pathway involves multiple molecular interactions, such as proteins degraded by the proteasome, peptide fragments transported into the endoplasmic reticulum by the transporter associated with antigen processing (TAP), and MHC Class I molecules binding to these peptides and presenting them on the cell surface for T-cell recognition (a process often referred to as peptide presentation or elution). MHC class I molecules are highly polymorphic (Radwan et al., 2020), and humans carry varying combinations of alleles, with the specific set differing across individuals. While datasets exist for various stages of this pathway, the most data-rich resources are those for peptide elution, binding affinity, and cleavage site prediction (Wohlwend et al., 2025; Hoof et al., 2009; Dorigatti et al., 2022). However, several challenges remain, such as imbalanced datasets, MHC polymorphism introducing variability in data points across alleles, and the multi-step nature of the pathway making immune response challenging to model *in-silico*. Previous work has typically focused on individual components, such as elution, binding, or cleavage, but in isolation. While some models achieve partial success, they often lack uncertainty estimates or measures of human tolerance, limiting their use in guiding protein design toward sequences with reduced epitopes. In this work, we hypothesize that, in the absence of large-scale labeled datasets distinguishing immunogenic from non-immunogenic sequences, a critical strategy is to design proteins with reduced immune response is through modeling the joint likelihood of all molecular interactions in the pathway.

Here, we use optogenetic proteins to test our platform (Maimon et al., 2018). Channelrhodopsins are seven-helix transmembrane microbial-derived proteins, that are widely used as molecular tools in neuroscience to control neurons, circuits, and behavior, yet their therapeutic applications have been limited by loss of expression and efficacy due to immune responses mediated by humoral immunity (Maimon et al., 2018). Proteins with complex functions, such as channelrhodopsins, require modeling of intricate conformational dynamics, photonic interactions, and gating transitions (channel opening and closing). These tasks remain beyond the capability of current machine-learning–based protein design pipelines where there is an absence of large training datasets. Traditionally, such challenges have been previously addressed through the construction of recombination libraries followed by screening for desired properties, but this requires a massive library synthesis and extensive characterization of thousands of mutants, creating a major bottleneck for therapeutic development (Mateljak et al., 2019).

In this work, we address protein redesign through a multi-faceted strategy. We perform recombination guided by protein language models, using blocks derived from closely related parent sequences, and show that as task complexity increases the success of trajectories decline, reflecting the expanding design space. We then apply GRPO with curriculum learning to the ESM-C model (Hayes et al., 2024; ESM Team, 2024), enabling the generation of samples with reduced elution and binding likelihoods and improved humanness scores across the five most frequent human alleles.

## 2 Related Work

The MHC class I antigen processing pathway has been modeled *in silico* in prior works, typically by focusing on individual components such as proteasomal cleavage or peptide–MHC presentation. Positive–unlabeled learning has been applied successfully to cleavage prediction (Dorigatti et al., 2022), while peptide presentation has been more extensively studied (Hoof et al., 2009; O’Donnell et al., 2020; Wohlwend et al., 2025), with recent models such as MUNIS achieving state-of-the-art performance by using pretrained protein language model to jointly encode the HLA sequence and the peptide sequence. Uncertainty estimation has also been explored in the context of peptide–MHC binding affinity prediction, for example in PUFFIN, a deep residual network–based framework (Zeng & Gifford, 2019). In this work, we employ evidential deep learning to provide unified uncertainty estimates across alleles, enabling sampling to be guided toward regions of higher certainty. Unlike earlier studies that align protein design outputs (e.g., from ProteinMPNN (Dauparas et al., 2022)) to predictors such as NetMHCpan (Hoof et al., 2009) using direct policy optimization, our framework leverages group relative policy optimization (GRPO). This enables us to jointly model multiple stages of the MHC Class I processing pathway without generating pairwise preference datasets.

There are multiple strategies to incorporate guided sampling into protein design models. ProteinMPNN conditions on structural context to generate sequences consistent with known folds, while RFdiffusion uses diffusion-based sampling with geometric constraints to produce structurally plausible backbones. SCHEMA Mateljak et al. (2019) provides a recombination-based strategy by identifying cut sites that minimize disruption of structural contacts, though it depends on accurate structural information. More recently, flow-matching approaches such as MOG-DFM (Chen et al., 2025) have been proposed for multi-objective peptide design, optimizing properties such as half-life, solubility, and binding affinity, though their effectiveness can vary across tasks. In contrast, we employ reinforcement learning with group-relative policy optimization (GRPO) (DeepSeek-AI et al., 2025), which enables direct optimization over non-differentiable properties of protein sequences and provides a fast framework for guiding sample generation.

## 3 Methods

### 3.1 Modeling Molecular Interactions of MHC Class I Pathway

#### 3.1.1 MHC Class I Elution and Binding Affinity

Evidential deep learning (EDL) models jointly learn the target along with estimates of both aleatoric and epistemic uncertainty (Sensoy et al., 2018; Amini et al., 2020). Given a dataset *D* of *N* paired training examples, 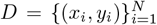, model learns a probability distribution over the likelihood parameters. For the MHC class I peptide elution task, we trained an evidential deep learning classifier with a Dirichlet prior, where given an HLA allele sequence and peptide, the model estimates evidence for a distribution over the binary outcome of elution versus non-elution.

##### Elution Prediction Classification

Let ***α*** ∈ ℝ^*K*^ denote the evidence parameter for *K* = 2 classes.

Define 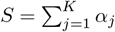. For a one-hot target vector **y**, the expected negative log-likelihood is

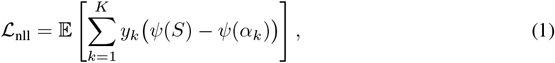

where *ψ*(·) is the digamma function. The full classifier loss combines this with a Kullback–Leibler regularizer:

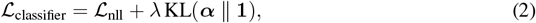

where *λ* is a weighting coefficient.

Epistemic uncertainty is then calculated by first normalizing the Dirichlet parameters into class probabilities. For a given class *k*, the predictive probability is defined as *p*_*k*_ = *α*_*k*_*/S*, where *α*_*k*_ represents the evidence assigned to class *k* and 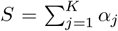 is the total evidence across all *K* classes. The predicted labe ŷ is then taken as the class with the highest probability,ŷ = arg max_*k*_ *p*_*k*_. The epistemic uncertainty is quantified as *u*_epistemic_ = *K/S*, which captures the model’s lack of knowledge: when the total evidence *S* is large, *u*_epistemic_ is small, indicating high confidence, whereas when *S* is small, *u*_epistemic_ grows large, indicating uncertainty about the prediction. The classifier was trained on a curated dataset of 651,237 unique peptides and evaluated on 41,725 positive HLA peptides and 208,625 randomly sampled decoys across 24 alleles from (Wohlwend et al., 2025).

##### Binding Affinity Prediction

To extend evidential deep learning to predict peptide–MHC binding affinity, we utilized the evidential regression framework introduced in Amini et al. (2020). This approach is based on the Normal–Inverse–Gamma (NIG) distribution, which serves as a conjugate prior over the mean and variance of a Gaussian likelihood function. The model outputs four hyperparameters (*γ*, ν, *α, β*) that parameterize the NIG distribution. The training objective combines two loss terms:

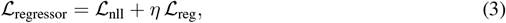

where the negative log-likelihood term is given by:

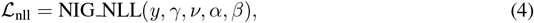

and the regularization term is:

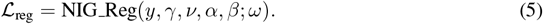

Here, *η* is the regularization coefficient. The evidential framework naturally decomposes uncertainty into epistemic and aleatoric components: 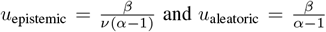. The regression model was trained on 182,203 peptide-MHC binding affinity datapoints and evaluated on 9,590 held-out test datapoints derived from the NetMHCpan dataset (Hoof et al., 2009).

#### 3.1.2 Proteasomal Cleavage

To predict proteasomal cleavage, we used datasets capturing both C-terminus and N-terminus sequence contexts for cleavage site recognition provided by (Dorigatti et al., 2022).The N-terminus dataset includes 1,285,659 samples with 229,163 positives, while the C-terminus dataset includes 1,277,344 samples with 222,181 positives. During policy optimization, only the C-terminus model was considered in the reward signal.

### 3.2 Human Tolerance Score

In the MHC class I pathway, peptides derived from both self and non-self antigens are displayed on the cell surface; however, self-derived (human) peptides typically do not elicit an immune response due to central tolerance. To account for this, we incorporate a weighting scheme that does not penalize peptides resembling human self-peptides when computing the reward during optimization. To calculate this tolerance, we constructed a position-specific scoring matrix (PSSM) using the human proteome, which is then used to evaluate the similarity of candidate 9-mer fragments to self-peptides. We focused on 9-mers, as they have been experimentally shown to be the most prevalent binders of MHC Class I alleles (Raghavan et al., 2025).

### 3.3 Protein Recombination through Protein Language Models

Protein function is often encoded in its evolutionary history, as captured by multiple sequence alignments (MSAs) of closely related protein sequences. Traditional methods such as SCHEMA (Mateljak et al., 2019) exploit this information by identifying optimal cut sites in protein sequences that minimize disruption to contacting residues and recombining blocks from homologous sequences to generate variants with desired functions. While SCHEMA introduces diversity through recombination, its restriction of recombination sites to avoid structural disruption limits its ability to achieve broad diversity while balancing function and immunogenicity. We illustrate this in Figure. 1b, where increasing the number of alleles to reduce epitopes results in fewer successful trajectories during MCMC sampling. In this framework, the Markov Chain Monte Carlo (MCMC) sampler replaces immunogenic blocks in a parent sequence with non-immunogenic blocks drawn from closely related parent sequences for a given set of alleles. Acceptance of replacements is determined by log-likelihood scores from the pretrained protein language model. While this approach leads to successful library designs for a limited set of alleles, the fraction of successful trajectories decreases as the number of distinct HLA alleles considered increases, as shown in Figure 1b. To address this scalability challenge, we employ GRPO to align language models and subsequently sample candidate sequences.

**Figure 1.**
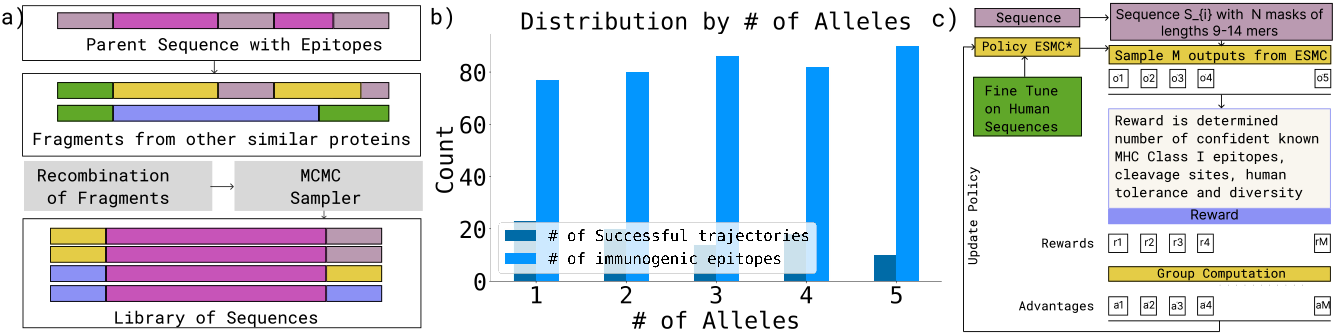
Framework for reducing immunogenic epitopes in protein design. (a) Recombination strategy showing how non-immunogenic segments from related sequences are swapped into parent proteins to generate chimeric libraries with fewer epitopes. (b) Distribution of outcomes as the number of distinct HLA alleles considered increases (x-axis), comparing number of successful trajectories (dark blue) out of 100 attempts to replace immunogenic epitopes (light blue) through the recombination strategy. (c) Schematic of the group relative policy optimization of ESM-C model for generating sequences with reduced MHC Class I epitopes.

#### Algorithm 1

MCMC Sampler For Chimeric Protein Library Generation to reduce T-cell Epitopes

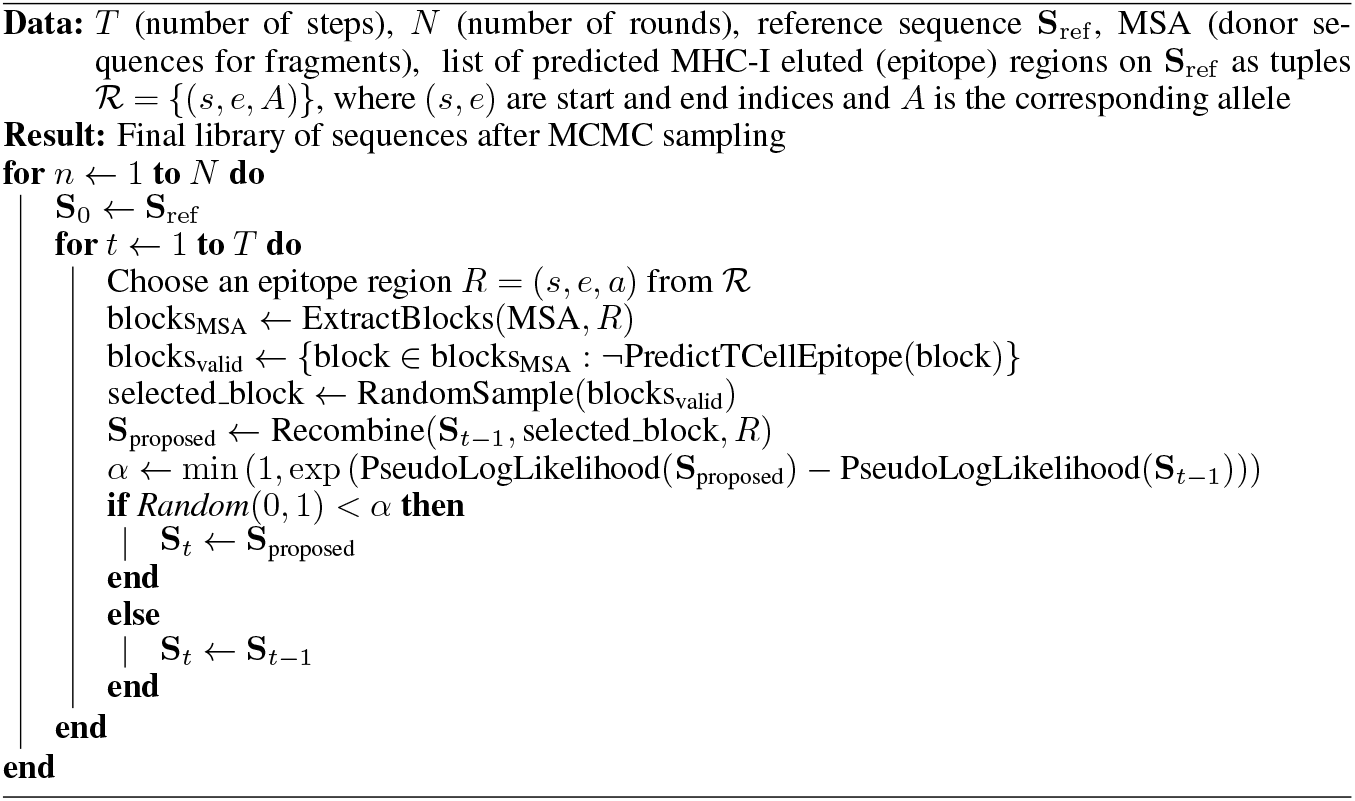

### 3.4 Supervised Fine-Tuning with Human Sequences

To enhance the model’s ability to generate sequences with reduced MHC Class I epitopes, instead of policy optimizing the base model directly, we fine tuned to produce tokens similar to human sequences through supervised fine tuning.

#### Algorithm 2

Supervised Fine-Tuning for Human peptide Proposal Generation

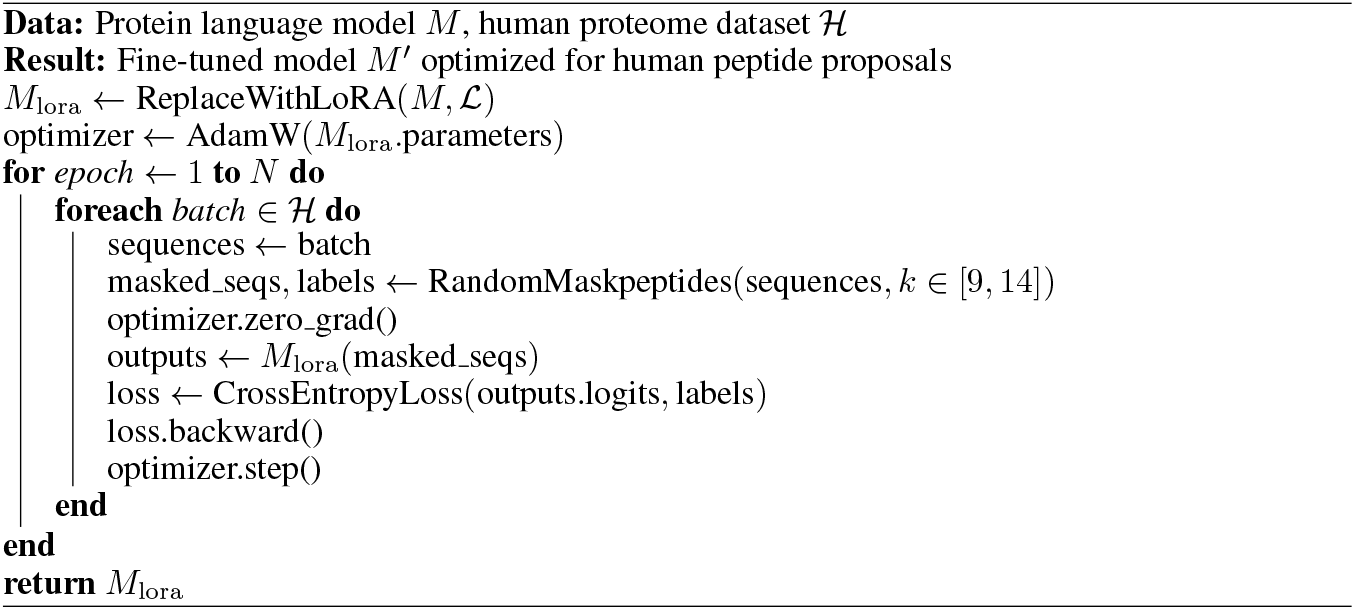

### 3.5 Group Relative Policy Optimization

We utilized GRPO with a handcrafted reward function to generate sequence proposals that exhibit reduced probabilities of elution and cleavage, higher human tolerance scores, and enforced diversity during sampling. To stabilize training, we incorporated a curriculum learning–based scheduler that progressively increases both the number of epitopes and the number of alleles considered, resulting in the GRPO variant CL-GRPO. Let 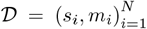 denote a dataset of protein sequences, where *s*_*i*_ ∈ Σ^*L*^ is a protein sequence of length *L* over the standard amino acid alphabet, and ⊂ *m*_*i*_ 1, 2, …, *L* represents the set of masked positions requiring optimization. To improve robustness, the reward also accounts for predictive uncertainty (penalizing overconfident but unreliable predictions) and similarity to human reference sequences (promoting tolerance). The policy model *M*_*θ*_ : Σ^*L*^ → ℝ^*L×*|Σ|^, parameterized by weights *θ*, is an ESM (Evolutionary Scale Modeling) protein language model enhanced with Low-Rank Adaptation (LoRA) for efficient fine-tuning. The reward function *R* : Σ^*L*^ × Σ^*L*^ → ℝ evaluates sequence quality by integrating MHC class I cleavage probability, peptide elution likelihood, and binding affinity, together with predictive uncertainty and similarity to human reference sequences.

#### Algorithm 3

Group Relative Policy Optimization with Curriculum Learning

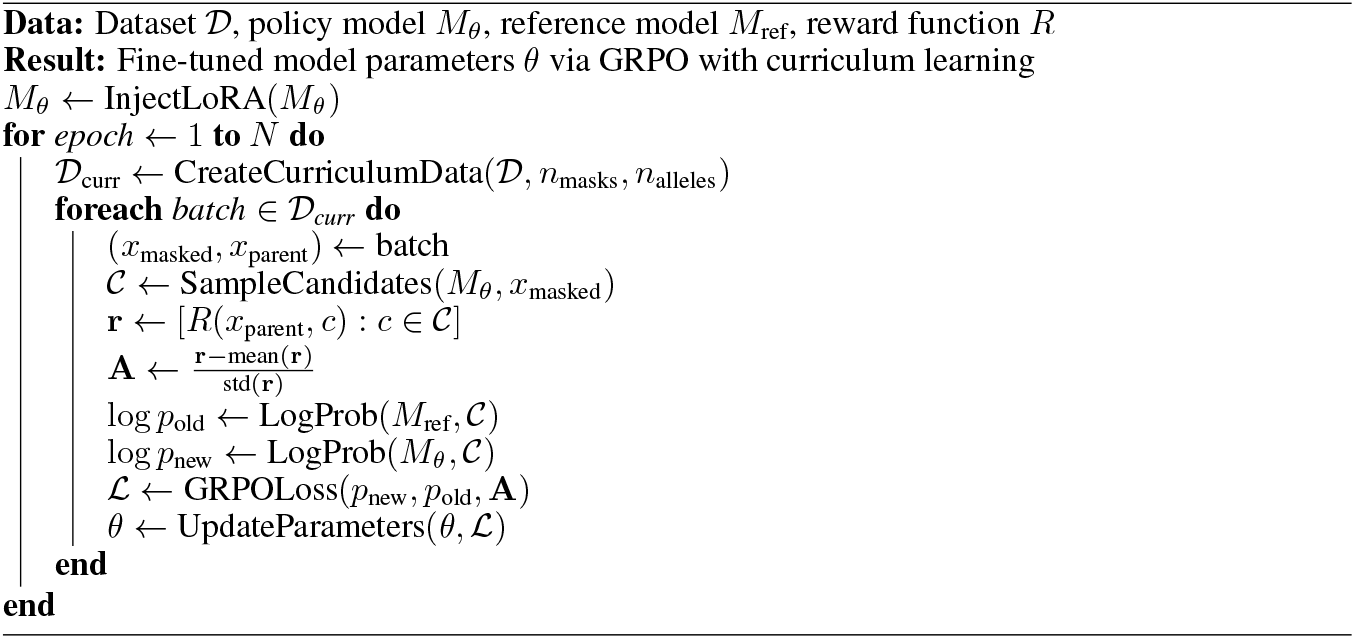

## 4 Results

### 4.1 MHC Class I Elution and Binding Affinity Prediction

We trained an evidential deep learning–based classifier using one-hot encoding for both the peptide and MHC sequence, which achieved strong predictive performance (AUROC = 0.963 on held-out test data; see Figure 2). The model also provides principled uncertainty estimates, which we use to weight the reward signal. As shown in Figure 2a, our model (PEARL-EL) performs competitively with existing benchmarks—while not the top performer, it reaches comparable accuracy with the advantage of calibrated uncertainty quantification and fast inference, making it well suited for use in reward scoring in policy optimization. Figure 2b and Figure 2c highlight performance variation across alleles and peptide lengths, respectively, showing that performance strongly depends on the allele of interest. This underscores that, during policy optimization, immunogenic peptides should not be weighted uniformly but instead adjusted according to allele-specific behavior. For the complementary binding affinity prediction task, we evaluated regression performance using RMSE and Pearson correlation. Binding affinities were transformed as

**Figure 2.**
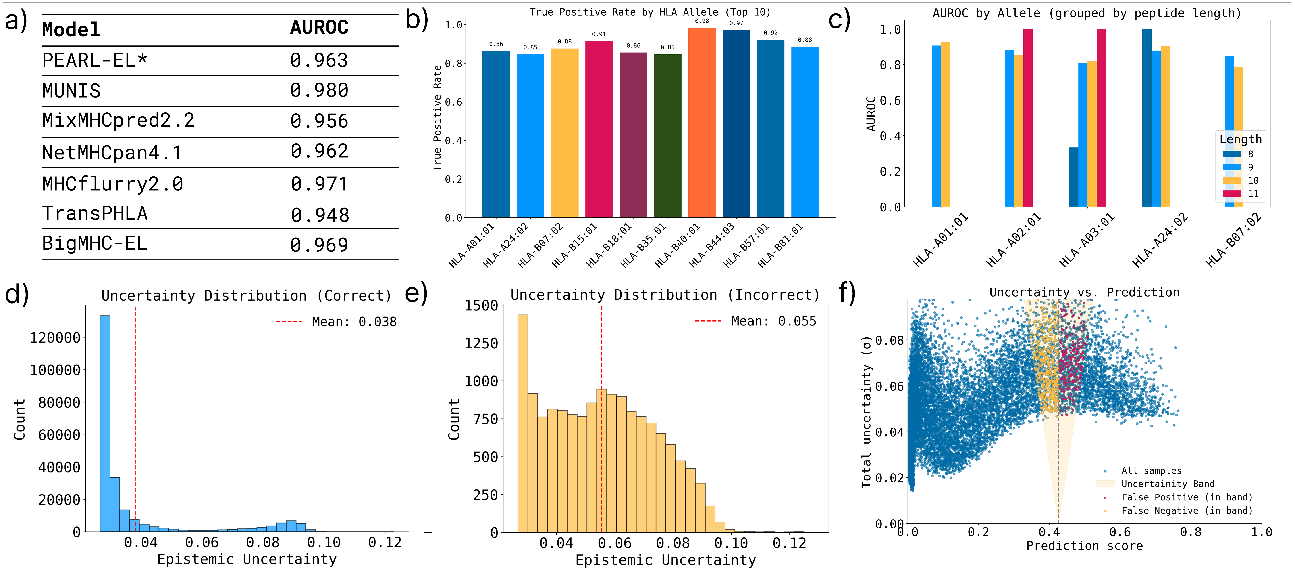
Evaluation of MHC Class I Elution and Binding Affinity Prediction (a) Comparison of PEARL-EL against existing benchmarks. (b-c) True positive rates across HLA alleles and peptide lengths demonstrate both predictive power and variability. (d–e) Epistemic uncertainty distributions of correctly classified and misclassified peptides respectively. (f) Uncertainty band showing the false positives and false negatives from binding affinity predictions.

**Figure 3.**
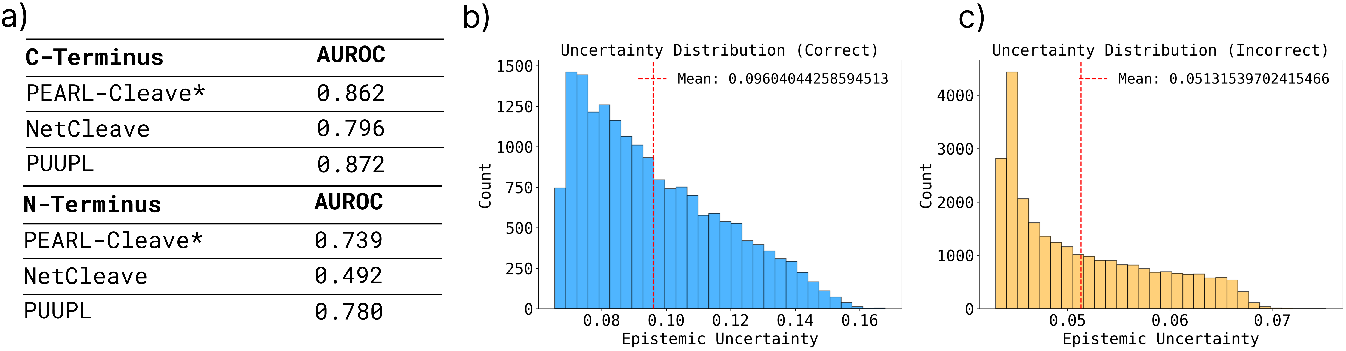
Evaluation of protease cleavage predictor.(a) AUROC for C-Terminus and N-Terminus predictions shows competitive accuracy of our approach. (b) Distribution of Epistemic uncertainities for predicting the correct label. (c) Distribution of Epistemic uncertainities for predicting the incorrect label.

**Figure 4.**
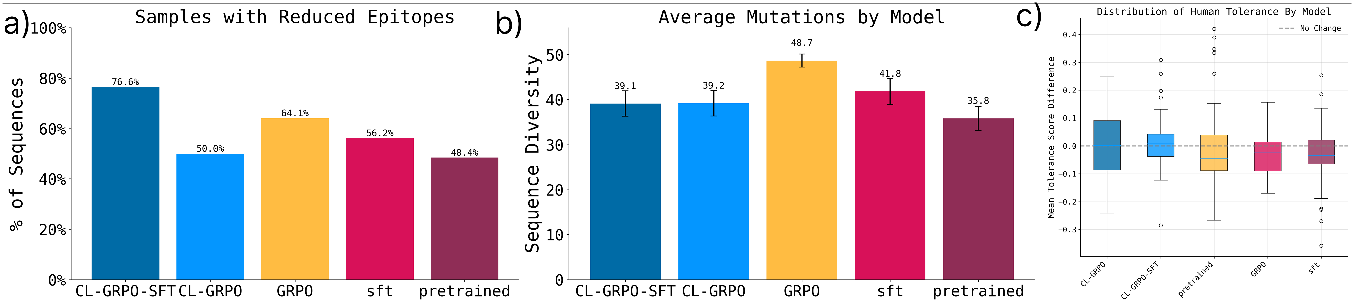
Reduction in MHC Class I epitopes through policy optimization. (a) Percentage of samples with reduced binder counts compared to other models across HLA-A02:01, HLA-A01:01, HLA-A03:01, HLA-A24:02, and HLA-B07:02, indicating where epitopes are effectively minimized. (b) Average number of mutations in the proposed sequences compared to the parent sequence. (c) Change in human tolerance score, which was incorporated into the reward function, showing that CL-GRPO and CL-GRPO-SFT perform strongly on this metric.

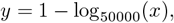

where *x* is the raw binding affinity in nM. This maps values into the range [0, 1], with affinities *x <* 500 nM considered binders. The evidential regressor not only produced uncertainty estimates but also discriminates between binders and non-binders, with per-allele AUROCs of 0.9259 for *HLAA01:01*, 0.8808–1.0 for *HLA-A02:01*, 0.8095–1.0 for *HLA-A03:01*, 0.8786–1.0 for *HLA-A24:02*, and 0.7875–0.8486 for *HLA-B07:02*. These results demonstrate that uncertainty-aware regression can explain the observed variability between “strong” and “weak” binders, a phenomenon also seen in NetMHCpan predictions as highlighted in Figure 2f.

### 4.2 Protease Cleavage Predictor

An important upstream step in antigen presentation is the proteasomal cleavage of the source protein into peptides. We trained a classifier to predict whether a given site in a protein sequence will be cleaved by the proteasome, i.e. yielding a peptide suitable for MHC I presentation.

For the cleavage prediction task, we trained two separate evidential classifiers for N-terminal and C-terminal contexts, following (Dorigatti et al., 2022). The C-terminal model achieved an AUROC of 0.862 on held-out data, outperforming NetCleave but below the PUUPL model. Using simple one-hot encoding, our approach enables fast inference for computing the reward while also providing uncertainty estimates. This is particularly valuable when integrating cleavage likelihood into policy optimization as a reward.

### 4.3 Reducing MHC Class I Epitopes Through Policy Optimization

We developed a reinforcement learning strategy to design protein sequences with reduced MHC class I epitopes. As a test case, we used a microbial-derived protein that lacks human sequence analogs and has been shown to elicit an immune response in vivo (Maimon et al., 2018). Specifically, we selected the well-studied channelrhodopsin C1C2 (a 350-residue light-gated ion channel, PDB ID: 3UG9) from the family of known optogenetic proteins.

To introduce mutations that minimize predicted MHC I presentation while maintaining protein stability and avoiding excessive sequence changes that could disrupt function, we defined a multiobjective reward function R that integrates outputs from our models: (1) a negative reward for peptides predicted to be presented by common HLA alleles using both elution and binding predictors, (2) a negative reward for peptides likely to be cleaved based on the protease model, and (3) a positive reward for higher tolerance similarity to human self-peptides. We also incorporated a diversity term to encourage exploration of sequence space, employing nucleus top-p sampling Holtzman et al. (2020) with temperature modifier to maintain sufficient diversity.

Training was performed with Group Relative Policy Optimization (GRPO), which normalizes and balances contributions from each reward component. To further stabilize learning, we implemented a curriculum schedule that gradually increased task difficulty—starting with a limited set of the most immunogenic regions and progressively allowing more mutations. Interestingly, we found that in some instances without this curriculum, GRPO could handle larger immunogenicity masks sampled at once and still achieve strong performance. Nonetheless, we believe curriculum learning will be especially valuable for scaling to settings with more alleles, where optimization complexity increases substantially.

Using the trained policy, we generated a set of 64 channelrhodopsin variants and evaluated their predicted epitope probabilities across multiple HLA class I alleles. Binder counts were predicted using NetMHCpan rather than our own models, providing an independent evaluation of epitope reduction. Figure 5a–e show the top eight CL-GRPO-SFT–designed variants for each of the five alleles, with separate plots for strong and weak binders. For almost all samples, we observed reductions in both strong and weak binders compared to the baseline channelrhodopsin. Figure 5f shows the distribution of AlphaFold confidence scores for the designed sequences, with a mean pLDDT of 84.78, indicating that structural integrity is broadly preserved.

**Figure 5.**
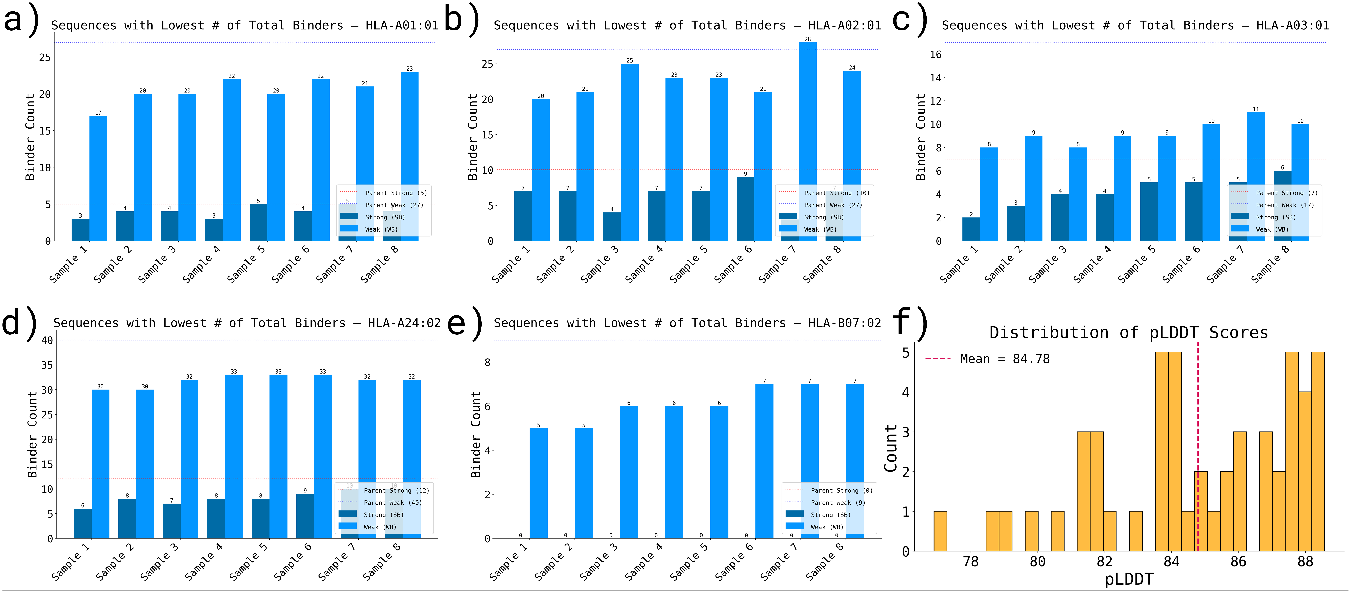
MHC Class I Epitope Prediction and Structural Evaluation. (a–e) Top eight sequences from the CL-GRPO-SFT model across HLA-A02:01, HLA-A01:01, HLA-A03:01, HLA-A24:02, and HLA-B07:02, showing consistent reduction in both strong and weak binders. Yellow bars indicate weak binders and blue bars indicate strong binders, with dotted red and yellow lines marking the corresponding counts for the parent reference sequence. (f) Distribution of average pLDDT scores from AlphaFold predictions for all optimized sequences.

Figure 5a–e present the top eight CL-GRPO-SFT–designed channelrhodopsin variants for each of the five HLA class I alleles, with separate plots showing predicted strong and weak binders as determined by NetMHCpan. Across nearly all samples, we observed consistent reductions in both strong and weak binders compared to the baseline sequence. Figure 5f shows the distribution of AlphaFold confidence scores for the designed variants, with a mean pLDDT of 84.78, indicating that structural integrity is broadly maintained. Further analyses and detailed comparisons are provided in the Appendix.

### 4.4 Discussion

In this work, we introduce PEARL (Protein Epitope Avoidance by Reinforcement Learning), an evidential deep learning–driven reinforcement learning framework for protein design aimed at reducing immunogenicity while preserving protein function and stability. The model incorporates elution, binding affinity, and cleavage scores as reward signals. Uncertainty estimates from these predictors are used to weight the rewards, ensuring that more confident predictions exert greater influence during policy optimization. By combining these predictors within a curriculum-guided reinforcement learning policy optimization, we generated optimized channelrhodopsin variants with substantially reduced predicted MHC class I epitopes. Compared to baseline protein language model, the policy optimized model consistently lowered the number of epitopes. Nonetheless, important challenges remain, particularly in scaling the framework and exploring deeper models to further enhance generalization. This work is limited to computational predictions, and experimental validation will be essential to confirm whether reduced epitope counts translate into lowered immune response *in vivo*. The framework can also be utilized for patient allele–specific redesign of proteins, and can further be expanded to include class II MHC presentation and other aspects of immune recognition such as T-cell receptor binding. More broadly, this approach has the potential to extend beyond channelrhodopsins to a wide range of therapeutic proteins.

## 4.5 Acknowledgements

S.S. acknowledge the Schmidt Science Fellows for their generous support through the postdoctoral fellowship. S.S also thanks the Schmidt Future’s Virtual Institute for Scientific Software (VISS program) and Johns Hopkins University’s Scientific Software Engineering Center (JHU SSEC) for computational support and valuable discussions. M.P. and J.J. acknowledge Eleven Eleven Foundation and NecSys at MIT Media Lab for their support and compute resources. ESB acknowledges HHMI, Lisa Yang, NIH R01DA029639, NIH R01MH122971, NSF 1848029, and John Doerr.

## 4.6 LLM Usage

A large language model was used to assist with text polishing, code templating, debugging, and generating scripts for plots in this paper, and LaTeX formatting for mathematical expressions and pseudocode.

