## Supplemental Figure for "Designing proteins with reduced T-cell epitopes through policy optimization"

### A Appendix

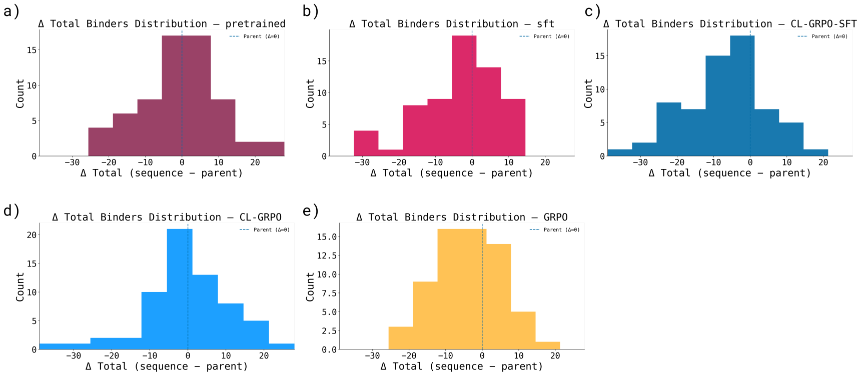


Figure 1: Distribution of Δ binder counts between parent and designed sequences. Panels a–e show the spread of (proposed sequence - parent sequence) binder counts for each allele. Negative values indicate a reduction in predicted binders relative to the parent sequence.

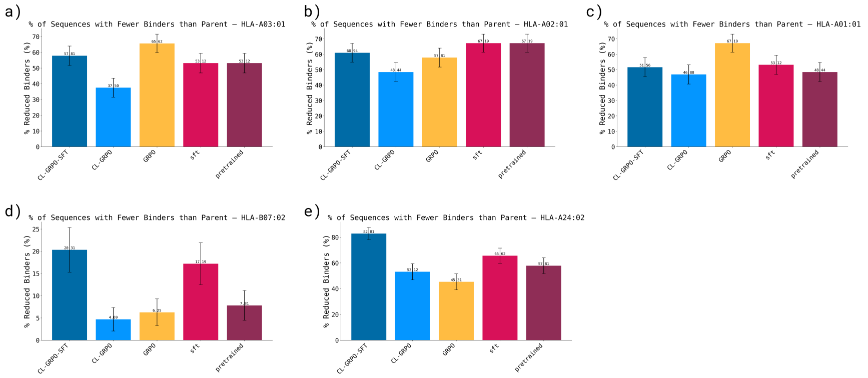


Figure 2: Number of samples with reduced binder counts compared to other models across each allele. Despite some variation in performance for individual alleles, CL-GRPO-SFT consistently achieves the greatest overall reduction in total predicted binders across all alleles.

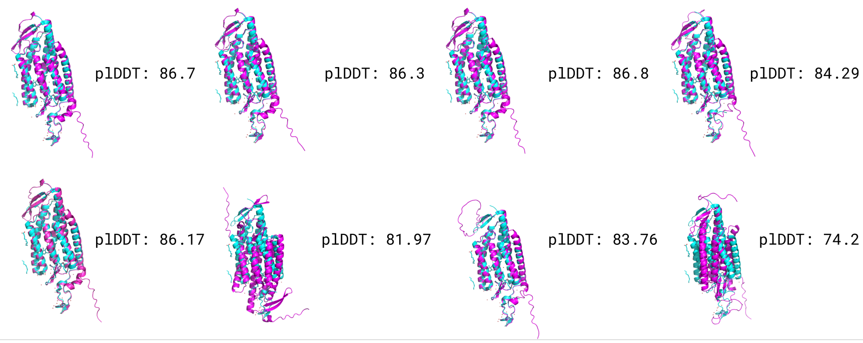


Figure 3: AlphaFold2 predicted structures of the CL-GRPO-SFT generated samples
